# Preserved efficacy and reduced toxicity with intermittent linezolid dosing in combination with bedaquiline and pretomanid in a murine TB model

**DOI:** 10.1101/2020.06.10.145334

**Authors:** Kristina M. Bigelow, Rokeya Tasneen, Yong S. Chang, Kelly E. Dooley, Eric L. Nuermberger

## Abstract

The novel regimen of bedaquiline, pretomanid, and linezolid (BPaL) is highly effective against drug resistant tuberculosis, but linezolid toxicities are frequent. We hypothesized that, for a similar total weekly cumulative dose, thrice-weekly administration of linezolid would preserve efficacy while reducing toxicity compared to daily dosing, in the context of the BPaL regimen.

Using C3HeB/FeJ and BALB/c mouse models of tuberculosis disease, thrice-weekly linezolid dosing was compared to daily dosing, with intermittent dosing introduced (a) from treatment initiation or (b) following an initial period of daily dosing. In all animals, BPa was dosed daily throughout treatment. Blood counts were used to assess hematologic toxicity. Following unexpected findings of apparent antagonism, we conducted additional experiments to investigate strain-to-strain differences in the contribution of linezolid to regimen efficacy comparing each 1- and 2-drug component to the BPaL regimen in BALB/c mice infected with *Mycobacterium tuberculosis* H37Rv or HN878.

Giving linezolid daily for 1-2 months achieved the greatest efficacy, but following that, results were similar if the drug was stopped, dosed thrice-weekly, or continued daily. Erythrocyte counts were lower with daily than thrice-weekly dosing. Linezolid had additive effects with BPa against *M. tuberculosis* H37Rv but antagonistic effects with BPa against *M. tuberculosis* HN878. However, overall efficacy of BPaL was high and similar against both strains.

Dosing linezolid daily for the first two months, then less frequently thereafter, may optimize its therapeutic margin. Linezolid’s contribution to BPaL regimens may depend on *M. tuberculosis* strain.

## INTRODUCTION

Tuberculosis (TB) is the leading cause of death by a single infectious agent worldwide. While treatment of drug-susceptible TB is relatively effective, multidrug-resistant (MDR) (resistant to at least rifampin and isoniazid) TB and extensively drug-resistant (XDR) (resistant to at least rifampin, isoniazid, fluoroquinolones and a second-line injectable agent) TB have had far lower treatment success rates at 56% and 39%, respectively (1). To improve MDR/XDR-TB treatment success and shorten the duration of treatment, new regimens comprised of dose-optimized new and repurposed anti-TB drugs are needed.

The Nix-TB trial recently showed that a novel, short-course oral regimen comprised of bedaquiline (B), pretomanid (Pa) and linezolid (L) given for 26 weeks produced favorable long-term outcomes in 98 (90%) of 109 patients with MDR/XDR-TB who had no other treatment options (2). Despite these promising efficacy results, the majority of participants in the trial had to temporarily discontinue and/or reduce the dose of linezolid, due to toxicities including myelosuppression, peripheral neuropathy, or optic neuropathy (2, 3). Strategies to mitigate these linezolid-induced toxicities without compromising the efficacy of BPaL would enhance the field implementation of the potentially transformative BPaL regimen by reducing the intensity of toxicity monitoring and the need for adjustments to regimen composition. However, despite recent guidelines ranking linezolid as a preferred agent for treatment of MDR/XDR-TB, there is not consensus on the optimal dosing strategy to balance efficacy and toxicity.

The narrow therapeutic window of linezolid and the long treatment durations required for TB therapy make it difficult to identify a single dose or dosing schedule that optimizes efficacy while minimizing toxicity (2–5). Both its activity against *Mycobacterium tuberculosis* (*M. tuberculosis*) and its treatment-limiting toxicities appear “time-dependent”. Hematological, and possibly neuropathic, toxicities are most closely associated with the proportion of the dosing interval for which concentrations exceed the IC_50_ for mitochondrial protein synthesis (T_>MPS50_) or its surrogate, C_min_ (6–12). Likewise, the activity of linezolid against actively replicating *M. tuberculosis*, both in an *in vitro* hollow fiber model of TB and in an acutely infected BALB/c mouse model of TB, is most closely associated with time above MIC (T_>MIC_) (6, 13). However, we recently found that when bacterial growth is suppressed by the mouse adaptive immune response or co-treatment with a clinically relevant dose of pretomanid, the bactericidal effect of linezolid becomes less dependent on T_>MIC_ (13), suggesting that, when combined with pretomanid and other strong companion drugs, the same weekly dose of linezolid may be dosed more intermittently to limit T_>MPS50_-driven toxicity without significant loss of efficacy. These findings led us to hypothesize that, in the context of the BPaL regimen, thrice-weekly administration of linezolid would have similar efficacy and less hematological toxicity than the same total weekly dose of linezolid administered daily. We further hypothesized that “front-loaded” regimens with daily linezolid dosing at a dose equivalent to 1200 mg daily in humans for the first 1-2 months prior to switching to thrice-weekly administration of the same dose (i.e., lowering both total weekly dose and dosing frequency) would be the optimal strategy for maximizing efficacy while minimizing toxicity.

We set out to test the above dosing strategy hypotheses in a murine model of TB, first using C3HeB/FeJ mice, which are prone to development of necrotizing granulomatous lung lesions that better represent the pathology of active TB in humans (14–21), and then using the high-dose aerosol infection model in BALB/c mice in which the BPaL regimen was discovered (22). An alternative dosing strategy of administering linezolid for only the first 2 months, then continuing with BPa only, which is being tested in the ZeNix trial (ClinicalTrials.gov Identifier: NCT03086486), was also assessed.

## RESULTS

### MICs

Linezolid MIC against both strains was 0.5 μg/mL by agar proportion, while median MIC by broth microdilution was 1 μg/mL against H37Rv and 0.5 μg/mL against HN878. Bedaquiline MIC against both strains was 0.03 μg/mL by agar proportion, while median MIC by broth microdilution was 0.25 μg/mL against H37Rv and 0.125 μg/mL against HN878. Pretomanid MIC by agar proportion was 0.06 μg/mL against H37Rv and 0.125 μg/mL against HN878, but MICs by broth microdilution were 0.125 and 0.06 μg/mL, respectively.

### Experiments 1 and 2: Evaluating linezolid dosing strategies in C3HeB/FeJ mice

#### Experiment 1: HN878 strain

We set out to test our dosing strategy hypotheses in C3HeB/FeJ mice infected with *M. tuberculosis* HN878, which, in our experience, consistently produces necrotizing granulomatous lung lesions in a larger proportion of mice than *M. tuberculosis* H37Rv. Control mice received BPa alone, while other mice received BPa plus linezolid at 45 mg/kg or 90 mg/kg daily (6 days per week [6/7]), to produce plasma AUC values comparable to 600 mg and 1200 mg, respectively, in humans (13). To test the hypothesis that more intermittent dosing of linezolid would be equivalent to daily dosing of the same total weekly dose, another group received BPa plus linezolid at 90 mg/kg thrice weekly (3/7). After 2 months of treatment, unexpectedly, mice receiving BPaL had higher median CFU counts than animals treated with BPa alone (Figure 1A) suggesting the possibility of antagonism by linezolid when added to the regimen. None of the CFU count differences between the BPa group and each individual BPaL group was statistically significant. However, when the BPaL arms were combined *post hoc* to increase the power to detect a difference and compared to BPa alone, BPa alone was significantly more active than BPa plus linezolid (p = 0.026). This result indicating antagonism by linezolid was surprising because previous studies in our high-dose aerosol infection model in BALB/c mice have consistently shown that the addition of linezolid increases the efficacy of BPa after 1-3 months of treatment (22). Those studies used *M. tuberculosis* H37Rv as the infecting strain, raising the possibility that this unexpected result could be related to differences in the *M. tuberculosis* strains or to differences in mouse species.

**Figure 1.**
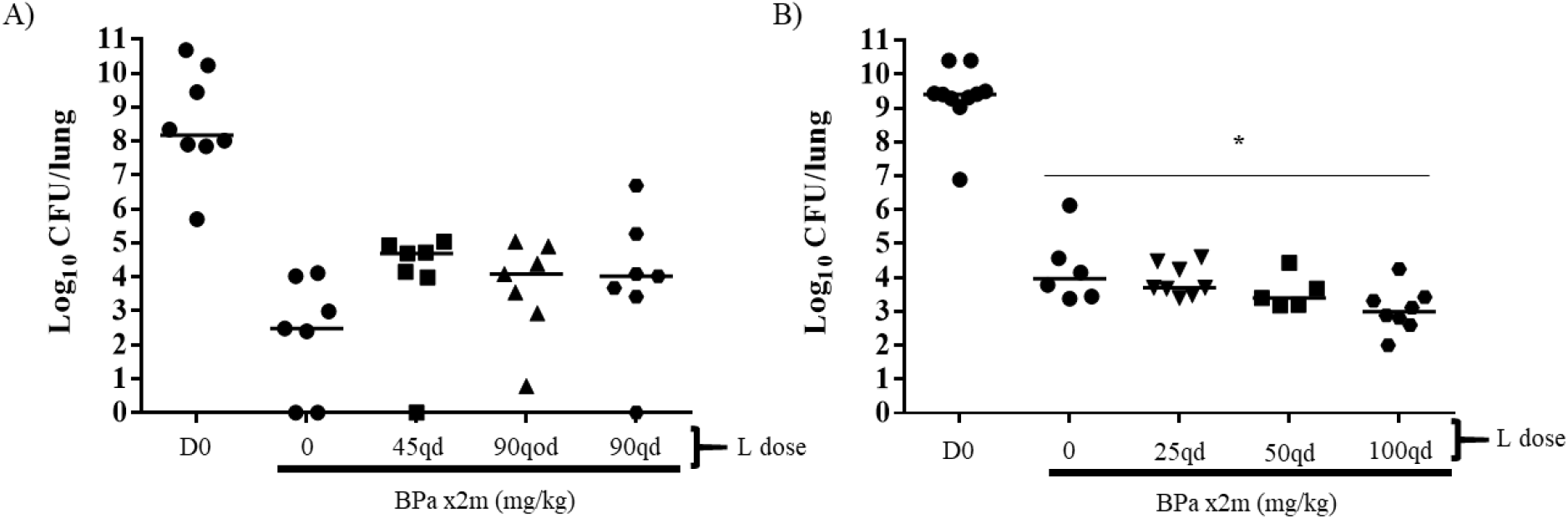
Lung CFU counts in C3HeB/FeJ mice after treatment with BPa with or without L at differing doses and dosing frequencies. A) Comparison of BPa with or without at L at 45 mg/kg qd, L 90 mg/kg qod, or L 90 mg/kg qd after 2 months of treatment in HN878-infected mice in Experiment 1. B) Comparison of BPa with or without at L at 25 mg/kg qd, 50 mg/kg qd, or 100 mg/kg qd after 1 month of treatment in H37Rv-infected mice in Experiment 2. Abbreviations: B, bedaquiline; Pa, pretomanid; L, linezolid; CFU, colony-forming unit; m, month; qd, once-daily dosing (5/7). Doses used: B 25 mg/kg qd, Pa 50 mg/kg (A) or 100 mg/kg qd (B). *, p-value<0.05. Dark horizontal bars indicate the median value for each group.

#### Experiment 2: H37Rv strain

When increasing doses of linezolid were added to BPa in the treatment of C3HeB/FeJ mice infected with the H37Rv strain of *M. tuberculosis* in **Experiment 2**, a dose-dependent increase in bactericidal activity was observed as the linezolid dose increased from 25 mg/kg, to 50 mg/kg and then to 100 mg/kg, and BPaL with linezolid dosed at 100 mg/kg was significantly more active that BPa (Figure 1B). After these results we moved to the simpler BALB/c mouse model to test both the original linezolid dosing strategy hypotheses and to determine whether the contribution of linezolid to the BPaL regimen was dependent on the *M. tuberculosis* strain.

### Experiment 3: Evaluating linezolid dosing strategies and bacterial strain-dependent antagonism of linezolid in BALB/c mice

In this experiment, we used the high-dose aerosol infection model in BALB/c mice infected with *M. tuberculosis* H37Rv or HN878. At the start of treatment (D0), mice infected with the H37Rv strain had a mean lung CFU count of 7.8 log10 while mice infected with the HN878 strain had approximately 6.5 log_10_ CFU, owing to a lower infectious dose of the latter strain. Mice infected with H37Rv received up to 3 months of treatment. BPa was given for the entire treatment duration. Controls received BPa only or BPa plus linezolid at 90 mg/kg once daily (5 days per week [5/7]) throughout. Test mice received BPa plus linezolid at 45 mg/kg (5/7) or 90 mg/kg (3/7), or linezolid was dosed once daily at 45 mg/kg or 90 mg/kg (5/7) for 1 or 2 months before being stopped or changed to 90 mg/kg (3/7). Additional controls received the standard drug-susceptible TB regimen of rifampin, isoniazid, pyrazinamide and ethambutol (RHZE).

As expected in mice infected with the H37Rv strain, the addition of daily linezolid to BPa increased the bactericidal activity of the regimen in a dose-dependent fashion (Table 1 and Fig. S1). Furthermore, BPa alone was significantly less bactericidal when compared to any linezolid-containing arm after 2 and 3 months of treatment (Table 1 and Fig. S2A-S2D). The same total dose of linezolid administered as 45 mg/kg (5/7) or 90 mg/kg (3/7) had similar additive bactericidal effects when combined with BPa, irrespective of whether the intermittent linezolid dosing began at the start of treatment or after 1 month of daily treatment with linezolid at 45 mg/kg. Likewise, no significant differences in the proportions of mice relapsing were observed when mice received linezolid at 90 mg/kg (3/7) instead of 45 mg/kg (5/7), confirming our hypothesis that, when administered with BPa, thrice weekly dosing of linezolid at 1200 mg would not compromise efficacy compared to linezolid 600 mg 5/7 (Table 1 and Fig. S2E and S2F) (13).

**Table 1.**
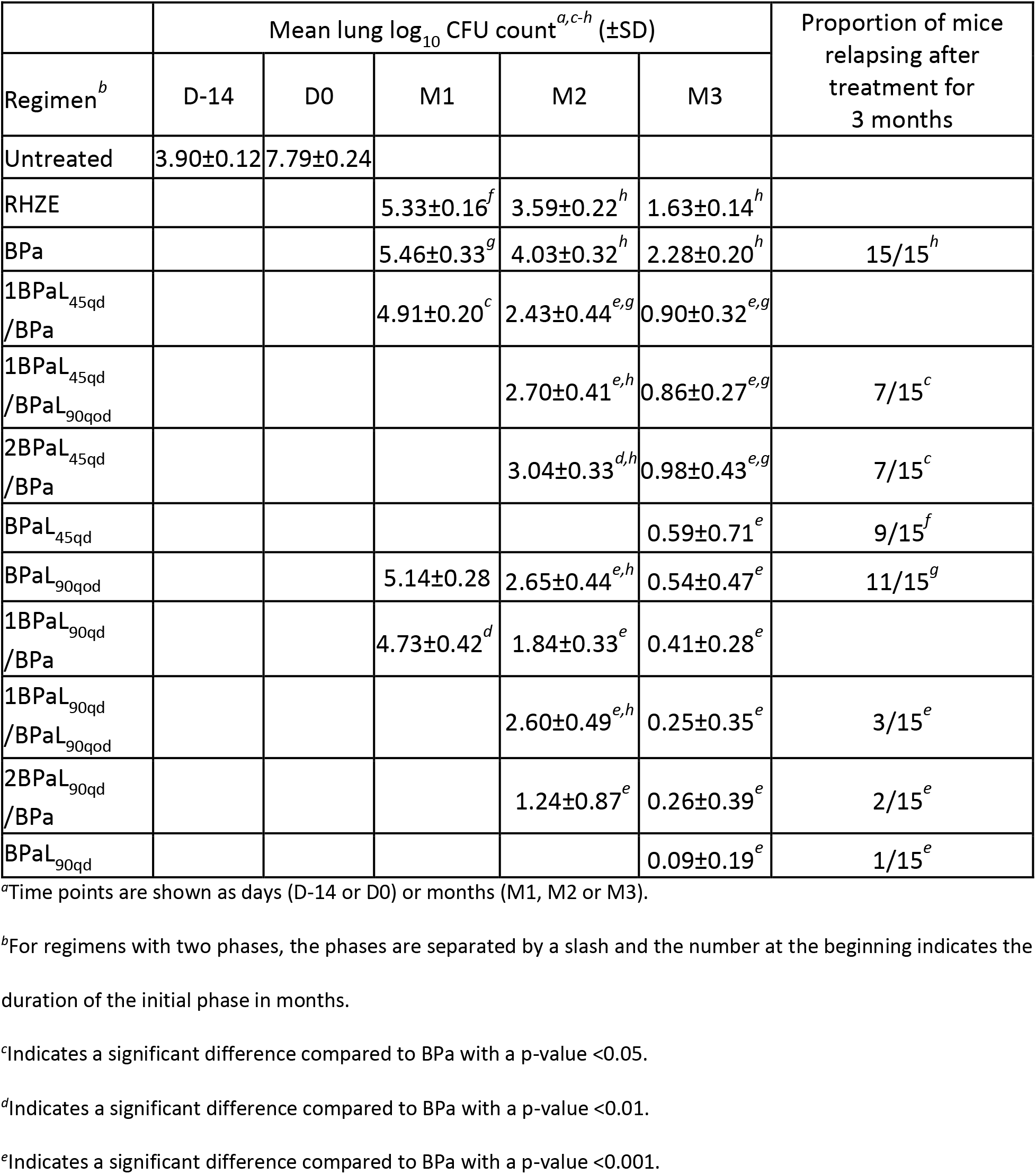
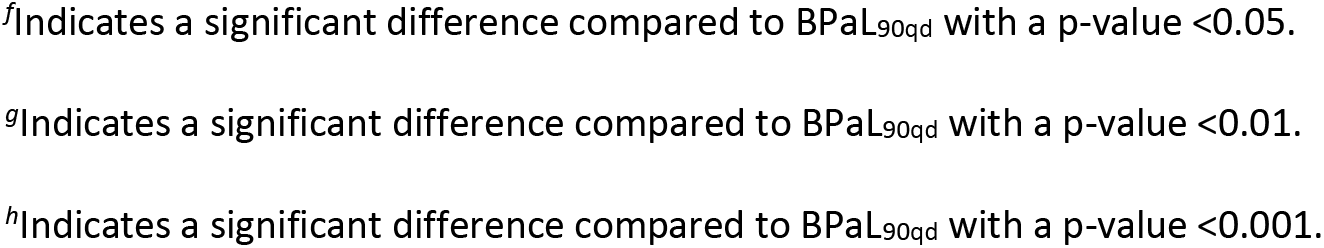
Lung CFU counts assessed during treatment and proportions of mice relapsing after treatment of BALB/c mice infected with *M. tuberculosis* H37Rv.

Regimens using linezolid 90 mg/kg (5/7) (the highest total weekly dose of any regimen), especially for the first 2 months, were the most efficacious (Table 1 and Fig. S2A-S2D). After 2 months of treatment, BPa plus linezolid 90 mg/kg (5/7) treatment was significantly more bactericidal than all other treatment arms except when linezolid 90 mg/kg (5/7) was discontinued at 1 month of treatment (Table 1 and Fig. S2A and S2B). After 3 months of treatment, BPa plus linezolid 90 (5/7) was significantly more bactericidal than all linezolid 45 mg/kg treatment arms except for when linezolid 45 mg/kg was given for the full duration of treatment (Table 1 and Fig. S2C and S2D).

Interestingly, giving the same total linezolid dose over 2 months by administering linezolid at 90 mg/kg (5/7) for just the first month resulted in lower CFU counts at month 2 than using linezolid at 45 mg/kg (5/7) or 90 mg/kg (3/7) for 2 months (Table 1 and Fig. S2A-S2D). An advantage of using linezolid at 90 mg/kg (5/7) was also observed in the proportions of mice relapsing after 3 months of treatment (Table 1 and Fig. S2E and S2F). Use of linezolid 90 mg/kg (5/7) for at least the first two months of treatment or linezolid 90 mg/kg (5/7) for 1 month followed by linezolid 90 mg/kg (3/7) was associated with relapse in only 1-3 out of 15 mice, compared to relapse proportions of 7-11 out of 15 mice when the linezolid dose was 45 mg/kg (5/7) and/or 90 mg/kg (3/7) (Table 1). The regimen using linezolid 90 mg/kg (5/7) throughout had statistically significantly fewer relapses compared to all groups having 9-11 relapses, even when adjusted for multiple comparisons, while the differences with groups having 7 relapses were only statistically significant before adjustment for multiple comparisons. By pairwise comparisons, the regimens using linezolid 90 mg/kg (5/7) for just the first 2 months or for the first month before switching to 90 mg/kg (3/7) produced significantly fewer relapses than the regimens using linezolid 45 mg/kg (5/7) or 90 mg/kg (3/7) throughout, although the difference was not quite statistically significant (p= 0.06) when comparing 1BPaL_90qd_/BPaL_90qod_ (3/15 relapses) to BPaL_45qd_ (9/15 relapses). Furthermore, 2 of the 3 mice relapsing after 3 months of the 1BPaL_90qd_/BPaL_90qod_ regimen had only a single CFU isolated (Fig. S2F). Taken together, these data support the hypothesis that initial dosing of linezolid at 90 mg/kg daily for the first 1-2 months results in superior efficacy compared to regimens with lower initial linezolid doses.

To assess the impact of linezolid dosing strategy on its hematological toxicity, whole blood was collected for complete blood counts (CBC) after 2 months of treatment in a cohort of uninfected mice treated in parallel with the infected mice. As previously observed (13), daily (5/7) dosing of linezolid at 45 or 90 mg/kg was associated with decreased red blood cell (RBC) indices (Figure 2A) compared to BPa-treated and/or untreated mice, whereas use of linezolid at 90 mg/kg (3/7) throughout or after an initial month of linezolid 90 mg/kg (5/7) was not (Figure 3A, 3B and 3C). These data suggest that decreasing linezolid dosing frequency, even after the first month of daily treatment, is associated with lower hematological toxicity. As in our previous study, platelet counts were not significantly affected (Figure 2D) (13).

**Figure 2.**
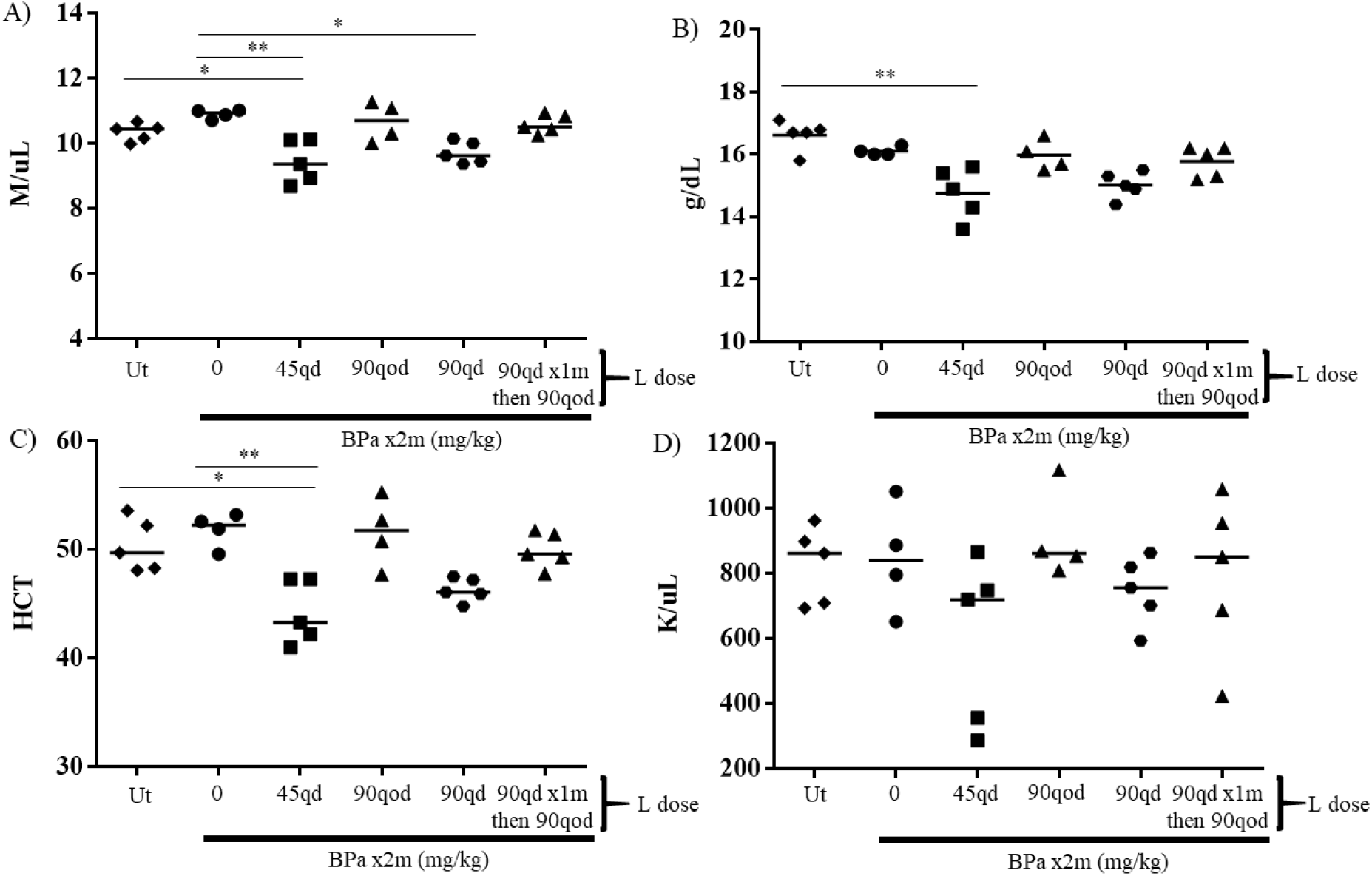
Results of complete blood counts performed in BALB/c mice after 2 months of treatment with BPa with or without L at differing dosing frequencies and doses in Experiment 3. A) RBC counts. B) Hemoglobin concentrations. C) Hematocrits. D) Platelet counts. Abbreviations: B, bedaquiline; Pa, pretomanid; L, linezolid; Ut, untreated; m, month; qd, once-daily dosing (5/7); qod, every other day dosing (3/7). Doses used: B 25 mg/kg qd, Pa 50 mg/kg, L as indicated. *, p- value<0.05; **, p-value<0.01;***,p-value<0.0001.

**Figure 3.**
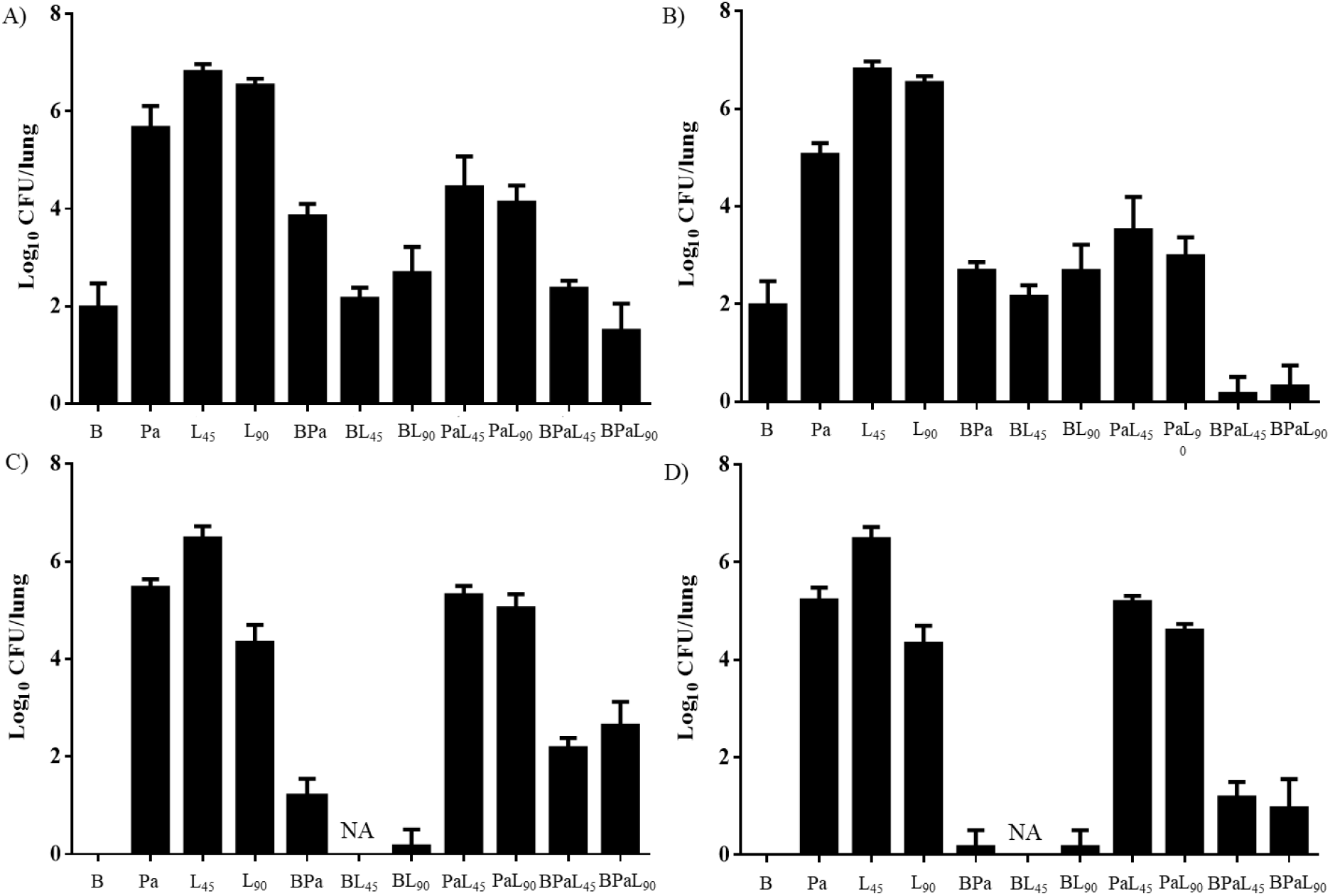
Lung CFU counts after 2 months of treatment with one, two and three drug combinations of B, Pa and L in BALB/c mice infected with *M. tuberculosis* strains H37Rv or HN878 in Experiment 4. A) Results for combinations using Pa 50 mg/kg qd in mice infected with *M. tuberculosis* H37Rv. B) Results for combinations using Pa 100 mg/kg qd in mice infected with *M. tuberculosis* H37Rv. C) Results for combinations using Pa 50 mg/kg qd in mice infected with *M. tuberculosis* HN878. D) Results for combinations using Pa 100 mg/kg qd in mice infected with *M. tuberculosis* HN878. Abbreviations: B, bedaquiline; Pa, pretomanid; L, linezolid; CFU, colony-forming unit; qd, once-daily dosing (5/7), NA, not available due to contamination of samples. Doses used: B 25 mg/kg qd, L 45 or 90 mg/kg qd, as indicated.

To confirm that the antagonism of linezolid on BPa observed in C3HeB/FeJ mice infected with the HN878 strain was bacterial strain-dependent, using a different mouse model, we gave BALB/c mice infected with *M. tuberculosis* HN878 BPa alone or in combination with linezolid given at 45 mg/kg (5/7), 90 mg/kg (5/7) or 90 mg/kg (3/7) for 2 months, or with linezolid at 90 mg/kg (5/7) for 1 month before being either stopped or switched to 90 mg/kg (3/7). Contrary to results from experiments using the H37Rv strain in BALB/c and C3HeB/FeJ mice, but consistent with results against the HN878 strain in C3HeB/FeJ mice, the addition of linezolid did not increase the activity of BPa against the HN878 strain (Table 2 and Fig. S3). Rather, linezolid antagonized BPa in what appeared to be a dose-dependent fashion. After 1 month of treatment, mean CFU counts were significantly higher among mice receiving BPa plus linezolid at 90 mg/kg (5/7) or every other day compared to those receiving BPa alone (Table 2 and Fig. S3A). After 2 months of treatment, many mice, including all mice receiving BPa alone, were culture-negative, but all groups receiving linezolid had at least one mouse with detectable CFU counts (Table 2 and Fig. S3B). H37Rv-infected mice treated with BPa plus linezolid 90 mg/kg (5/7) for 2 months experienced a reduction in bacterial burden of approximately 6.5 log_10_ CFU (Table 1), compared to a decrease of 6.2 log_10_ CFU with the same treatment in HN878-infected mice (Table 2). This suggested that, despite the antagonistic effect of linezolid on BPa in HN878-infected mice, the BPaL regimen remained highly effective.

**Table 2.**
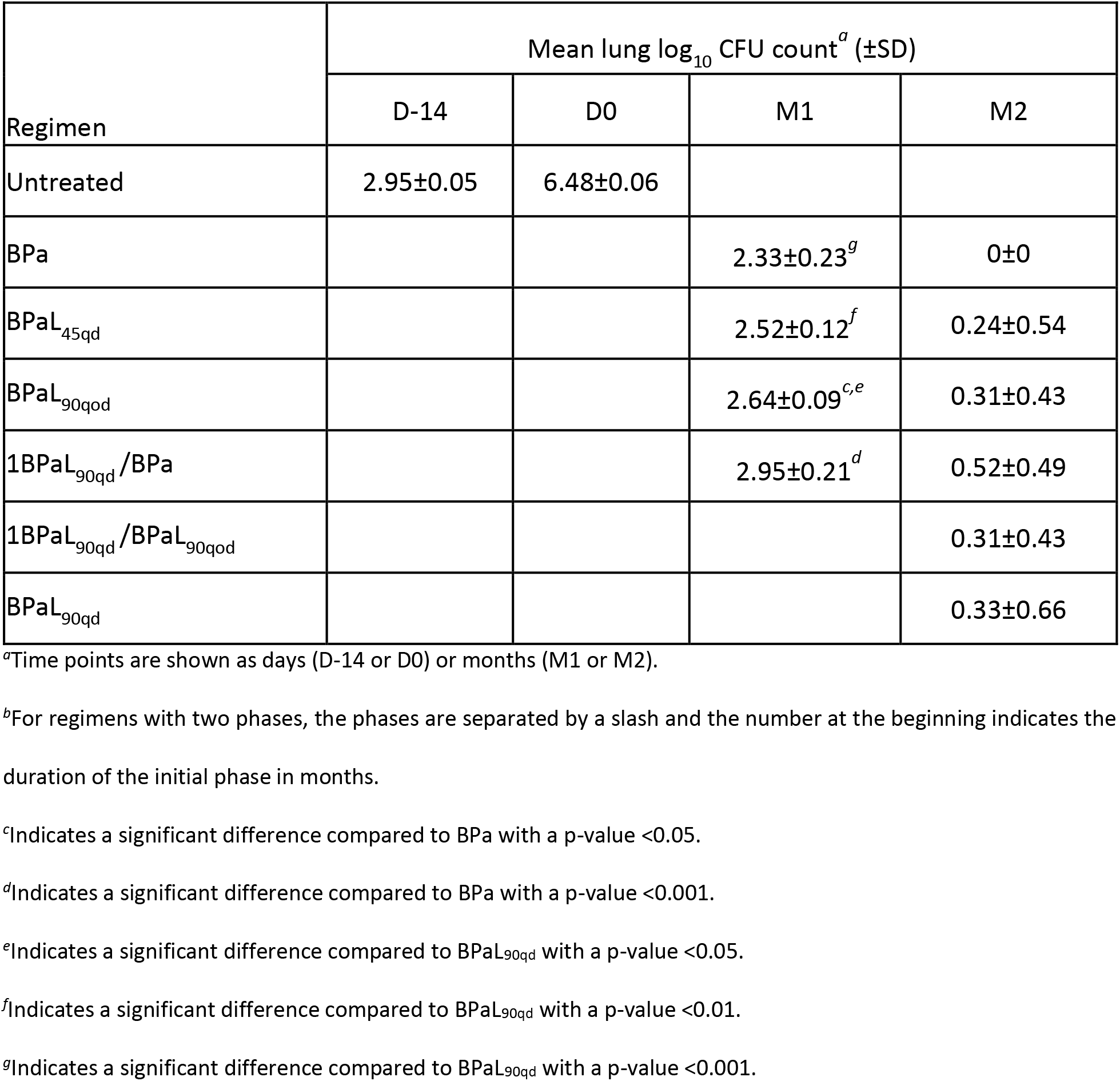
Lung CFU counts assessed during treatment and proportions of mice relapsing after treatment of BALB/c mice infected with *M. tuberculosis* HN878.

### Experiment 4: Evaluating interactions between drug components in the BPaL regimen and the influence of bacterial strain in BALB/c mice

To better understand the interactions between the component drugs and their contribution to the BPaL regimen, including the apparent bacterial strain-dependent antagonism of linezolid, BALB/c mice infected with either H37Rv or HN878 were treated with 1-, 2-, or 3-drug combinations of bedaquiline, pretomanid and linezolid (5/7) for 2 months. Pretomanid was dosed at 50 mg/kg or 100 mg/kg to achieve either a weekly AUC (50 mg/kg) or a T_>MIC_ (100 mg/kg) similar to that observed in patients taking 200 mg daily (23). Linezolid was dosed at either 45 mg/kg or 90 mg/kg. On D0 mice infected with H37Rv had mean lung CFU counts of 7.3 log_10_ CFU and HN878-infected mice had 8.7 log_10_ CFU. In mice infected with H37Rv, monotherapy with bedaquiline was more bactericidal than pretomanid or linezolid, and both pretomanid and linezolid had dose-dependent effects (Figure 3A and 3B). The addition of pretomanid at 50 mg/kg to bedaquiline alone decreased bedaquiline activity, while the addition of pretomanid at 100 mg/kg or either dose of linezolid had no significant effect. However, the addition of either dose of linezolid to pretomanid alone or to BPa resulted in additive effects, irrespective of the pretomanid dose, although only when the pretomanid dose was 100 mg/kg was the 3-drug combination more active than bedaquiline alone (Figure 3A and 3B).

The activity of bedaquiline alone was greater in mice infected with HN878 than those infected with H37Rv, rendering all HN878-infected mice culture-negative after 2 months of treatment (Figure 3C and 3D). As observed with H37Rv, addition of pretomanid at 50 mg/kg antagonized bedaquiline, while addition of pretomanid at 100 mg/kg or linezolid at 90 mg/kg had no significant effect. Samples from mice receiving bedaquiline plus linezolid at 45 mg/kg were contaminated during plating and could not be assessed. In stark contrast to results with H37Rv infection, no combination of pretomanid and linezolid was additive against HN878 and the addition of either linezolid dose to BPa was clearly antagonistic irrespective of the pretomanid dose. However, despite the differing contributions and interactions of the 3 component drugs, the full 3-drug BPaL regimen performed quite similarly against both *M. tuberculosis* strains.

To determine the effect of differing drug contributions to the selection of drug-resistant mutants, we enumerated CFU isolated on bedaquiline-containing and pretomanid-containing plates after 2 months of treatment (Fig. S4, S5 and Table S1). Consistent with prior observations (24), bedaquiline resistance (defined as ≥ 1% of total CFU) was observed in 2 of 5 H37Rv-infected mice treated with bedaquiline alone. Addition of linezolid at 45 mg/kg or pretomanid at 50-100 mg/kg prevented growth on bedaquiline-containing plates except for a single CFU on bedaquiline-containing plates for 2 mice or 1 mouse, respectively. Addition of both pretomanid and linezolid prevented recovery of any CFU on bedaquiline plates. Interestingly, no CFU were recovered on bedaquiline plates from any mouse infected with HN878 except for one mouse receiving BPaL with pretomanid 50 mg/kg and linezolid 90 mg/kg, the most antagonistic combination. Also consistent with prior observations, pretomanid resistance was selected in all mice given pretomanid alone (25), although 2 of 5 H37Rv-infected mice given pretomanid 50 mg/kg harbored only 0.02 and 0.1% resistant CFU. Addition of bedaquiline with or without linezolid prevented recovery of CFU on pretomanid plates except for a single mouse receiving bedaquiline and pretomanid at 100 mg/kg with 2 CFU on pretomanid plates. However, addition of linezolid only to pretomanid suppressed pretomanid resistance more effectively in HN878-infected mice (resistance in only 3 of 20 mice receiving any PaL combination) compared to H37Rv-infected mice (resistance in 13 of 20) (p= 0.003) despite greater additive effects of the combination against the susceptible bacterial population in H37Rv-infected mice (Fig. S4, S5 and Table S1).

Recently Pang et al. found antagonism between linezolid and bedaquiline in 13 out of 20 Chinese XDR-TB isolates using an *in vitro* checkerboard assay (26). To determine whether such antagonism might also occur in our HN878 strain, a member of the East Asian lineage, we performed 3 *in vitro* checkerboard assays to assess the interaction of linezolid and bedaquiline against both the H37Rv and HN878 strains. The first study used turbidity as the readout. Against the H37Rv strain, fractional inhibitory concentration index (FICI) values ranged from 1.06 to 1.5. Against the HN878 strain, the FICI range was 1 to 1.25. The next 2 studies used the Alamar blue assay, as in the study by Pang et al (26). Against the H37Rv strain, the FICI values were 0.56 to 1.25. Against the HN878 strain, the FICI values ranged from 0.75 to 2.25. All of these values fall in the “indifferent” or “no interaction” range (>0.5-4) (27). Therefore we did not identify antagonism between bedaquiline and linezolid against the H37Rv nor the HN878 strain in our *in vitro* checkerboard analysis. Notably, across these studies, the bedquiline MIC averaged 1 or 2 dilutions lower against the HN878 strain compared to the H37Rv strain, consistent with the greater efficacy of bedaquiline observed against HN878 compared to H37Rv *in vivo*.

## DISCUSSION

The potent sterilizing activity of the BPaL regimen and the contribution of each component drug was first discovered using a mouse model of TB (28). In the subsequent Nix-TB clinical trial, BPaL administered for 6 months resulted in an unprecedented 90% treatment success rate in therapeutically destitute patients with MDR/XDR-TB (2). Despite its promising efficacy as a short-course oral therapy for MDR/XDR-TB, use of the BPaL regimen is complicated by the dose- and duration-dependent toxicity of linezolid, which necessitated a linezolid holiday and/or dose reduction in most participants in the trial (2). No consensus exists on the optimal dosing strategy for linezolid to balance its efficacy and toxicity in TB treatment, whether in the context of the BPaL regimen or in other regimens (29). In the Nix-TB trial, linezolid was initiated at a daily dose of 1200 mg, but the dose was commonly reduced to 600 mg daily to manage toxicity, and linezolid was ultimately discontinued before the end of treatment in approximately one-third of participants. At least 2 clinical trials are underway to test different linezolid dosing strategies in the BPaL regimen. The TB-PRACTECAL trial (ClinicalTrials.gov Identifier: NCT02589782) is studying linezolid 600 mg daily for 16 weeks, followed by 300 mg daily or 600 mg every other day for the remaining 8 weeks of treatment. The ZeNix trial (ClinicalTrials.gov Identifier: NCT03086486) is studying linezolid at either 1200 mg or 600 mg daily, with both doses administered for either the entire 26 weeks or the first 9 weeks only.

Previous pre-clinical studies and limited clinical data have suggested alternative dosing strategies for linezolid. Summarizing results from *in vitro* hollow fiber system models of TB (6, 30, 31) and from our BALB/c mouse models, we recently proposed that, while linezolid efficacy is driven by T_>MIC_ when *M. tuberculosis* is growing exponentially in the absence of constraints, its efficacy is not as dependent on T_>MIC_ when *M. tuberculosis* multiplication is restricted by the host immune response or by combining with strong companion drugs (13). This led us to hypothesize that, when linezolid is combined with BPa, linezolid could be dosed at 1200 mg thrice weekly rather than 600 mg daily without loss of efficacy, provided that the same or similar total weekly dose is administered (13). By reducing the T_>MPS50_ (32), this thrice-weekly dosing strategy would be expected to reduce the risk of MPS-driven toxicity. However, because linezolid 1200 mg once daily is likely to be more bactericidal than linezolid 600 mg once daily or 1200 mg thrice weekly at the outset of treatment when *M. tuberculosis* replication is higher and before bedaquiline and pretomanid have reached steady state (6, 22, 32, 33), we also hypothesized that “front-loading” the regimen with linezolid at 1200 mg once daily before changing to 1200 mg thrice weekly after 1-2 months would be the most effective dosing strategy by increasing efficacy without significantly increasing the risk of toxicity compared to thrice-weekly linezolid throughout. A small case series suggesting mitigation of linezolid toxicity upon switching from 800 mg daily to 1200 mg thrice weekly further supports the front-loading concept (34).

The studies in the BALB/c mouse infection model reported here confirm these dosing strategy hypotheses. Replacing the 45 mg/kg daily dose of linezolid with 90 mg/kg thrice weekly did not compromise the bactericidal or sterilizing activity of the BPaL regimen and prevented reductions in RBC indices. As expected, replacing the 90 mg/kg daily dose of linezolid with 90 mg/kg thrice weekly was associated with reduced bactericidal and sterilizing activity, although it did again prevent reductions in RBC indices. Importantly, front loading the linezolid dosing with 90 mg/kg daily before switching to 90 mg/kg thrice weekly was more effective than linezolid 90 mg/kg thrice weekly throughout and was as effective as 90 mg/kg daily throughout, albeit with lesser effects on RBC indices than the last regimen. It is noteworthy that simply stopping linezolid after the first 2 months was as effective as continuing linezolid. This strategy of stopping linezolid at 2 months was initially suggested by a prior study in mice (22) and is currently being evaluated in the ZeNix trial. Our studies therefore provide additional evidence that stopping linezolid at 9 weeks in the ZeNix trial will most likely produce results similar to treating with linezolid for the entire treatment duration, while also minimizing toxicity since linezolid-induced toxicities are increasingly frequent after 2 months of treatment. However, continuing linezolid at a safer but still active dose may have advantages, particularly for protecting against the emergence of resistance to bedaquiline and/or pretomanid. Out of the 3 drugs in the BPaL regimen, linezolid has the highest barrier to resistance (35, 36). Indeed, prolonged administration of linezolid as functional monotherapy in XDR-TB patients was associated with selection of linezolid-resistant bacteria in only a minority of participants (3, 5). This suggests that continuing linezolid after the first 2 months could reduce the risk of emergent resistance to bedaquiline or pretomanid during treatment, an effect that the ZeNix trial may not be sufficiently powered to detect. With increasing evidence of baseline and acquired resistance to bedaquiline or nitroimidazole drugs among MDR/XDR-TB isolates (37–41), including acquired bedaquiline resistance in 1 of 105 participants treated with BPaL in a per protocol analysis, it may be prudent to keep linezolid in the regimen for the entire treatment duration but to change the dosing frequency after 1-2 months to thrice weekly to maintain efficacy but minimize toxicity.

We were surprised to find a decrease in efficacy when linezolid was added to BPa to treat infection with *M. tuberculosis* strain HN878 as opposed to the increase in efficacy observed when linezolid was added to BPa against H37Rv strain infection. Through our detailed study of the 1-, 2- and 3-drug combinations in the BPaL regimen, we found that the drugs interacted in and contributed to the regimen in different ways. Specifically, both bedaquiline and linezolid were significantly more active as monotherapy against the HN878 strain compared to the H37Rv strain, a finding that may be at least partly explained by the higher MICs in 7H9 media against the latter strain. A recent *in vitro* checkerboard study from Beijing reported antagonism between bedaquiline and linezolid against the majority of clinical XDR-TB isolates tested (26). However, we did not find such antagonism or evidence that the 2-drug pair of bedaquiline and linezolid interacted differently against H37Rv and HN878 in our *in vitro* checkerboard assay or in mice. Although pretomanid monotherapy did not exhibit differential effects, the interaction of linezolid with pretomanid was quite different between the two strains in BALB/c mice in that the combination of PaL was more active than pretomanid or linezolid alone against the H37Rv strain but not against the HN878 strain. This interaction of pretomanid and linezolid is what appeared to predict whether addition of linezolid to BPa would increase or decrease the efficacy. These results demonstrating a bacterial strain-dependent contribution of linezolid to the BPaL regimen raise the question of whether the observed differences are attributable to lineage-specific differences such that differential contribution of linezolid or differential activity of the regimen overall could be observed in geographic regions with high prevalence of isolates from the Beijing subfamily and/or East Asian lineage to which the HN878 strain belongs. Although it was out of scope for the current report, we plan to investigate whether similar differential contributions of linezolid in the regimen in mice occur with additional clinical isolates from the Euro-American and East Asian lineages. Another consideration is that the less favorable interaction of bedaquiline and/or pretomanid with linezolid is only observed at higher multiples of bactericidal exposures that were only reached against the HN878 strain owing to its greater susceptibility to bedaquiline and linezold. Such a mechanism may not be apparent in in vitro checkerboard studies based on growth inhibition but may be explored further through in vitro time-kill studies. In the end, it is important to note that, despite the strain-dependent differential contribution of linezolid to the regimen, the overall efficacy of the BPaL regimen was quite similar in BALB/c mice, irrespective of the infecting strain, because of the superior efficacy of bedaquiline against the HN878 strain. Moreover, despite not additive activity with pretomanid against the HN878 strain, linezolid was at least as effective at reducing selection of pretomanid-resistant mutants against this strain.

Our study has limitations. The pathological hallmarks of human pulmonary TB are caseating lung lesions and cavities. Although it provided the first evidence of the exceptional sterilizing activity of the BPaL regimen (22), the BALB/c mouse TB model used here produces only cellular lung lesions, and virtually all infecting bacilli reside intracellularly. We endeavored to test our dosing strategy hypotheses in a C3HeB/FeJ mouse TB model, which exhibits caseating lung lesions that may enable better representation of drug distribution into such lesions and activity against large extracellular bacterial populations in caseum (18, 42), but the unanticipated antagonism observed when L was added to BPa against the infecting HN878 isolate confounded our results. We successfully tested our dosing strategy hypotheses in the simpler BALB/c mouse model. Our intriguing results showing differential strain-dependent contributions of linezolid to the BPaL regimen are clearly limited by the use of only two strains. Further studies of additional strains from the East-Asian and Euro-American lineages will be necessary to determine whether these differences are lineage-dependent. Lastly, while we hypothesize that there may be clinical benefit in retaining linezolid beyond two months of treatment to prevent resistance, that hypothesis was not tested in these experiments.

## METHODS

### Mycobacterial strains

Experiments were conducted using *M. tuberculosis* H37Rv or HN878, as indicated. Unless otherwise noted, cultures were grown in Middlebrook 7H9 broth supplemented with 10% oleic acid-albumin-dextrose-catalase (OADC; Difco, Detroit, MI) and 0.05% Tween 80 (Sigma, St. Louis, MO). MICs were determined head-to-head against each strain using both the agar proportion method on Middlebrook 7H11 agar (performed once) and the microbroth dilution method in Middlebrook 7H9 medium without Tween 80 (Fisher, Pittsburgh, PA) (performed thrice for bedaquiline and linezolid, once for pretomanid).

### Aerosol infection model using BALB/c mice

All animal procedures were approved by the Animal Care and Use Committee of Johns Hopkins University. Six-week-old female BALB/c mice were used. Using an inhalation exposure system (Glas-Col, Terre Haute, IN) mice were infected with a log-phase culture of *M*. *tuberculosis* H37Rv or HN878 with an optical density at 600 nm of approximately 1.0 for a target dose of approximately 4 log_10_ CFU implanted in the lungs (22). After aerosol infection, mice were randomized into treatment groups (5 mice per treatment arm per time point during treatment and 15 mice per arm for relapse assessment, when evaluated). Untreated mice were sacrificed (i) the day after infection to quantify CFU implanted in the lungs, (ii) at initiation of treatment to determine pretreatment CFU counts, and (iii) 28 days post-infection to count CFU in untreated controls. Dosing (D0) started 14 days after infection.

### Aerosol infection model using C3HeB/FeJ mice

#### HN878 strain

Eight-week-old female C3HeB/FeJ mice were infected in a similar manner to BALB/c mice, except that lower infectious doses were used. A 1:400 dilution of an *M. tuberculosis* HN878 culture grown for 5 days in 7H9 media without Tween 80 was passed through a 25 gauge needle 3 times prior to aerosol infection. The infection was repeated 4 weeks later using the same protocol. After infection, body weight was monitored as an indication of disease progression. One cohort of mice with accelerated weight loss (defined as body weight ≤25g) started treatment 1 week after the second infection, after being randomized (4 mice per arm) into treatment arms. The remaining mice were held for an additional 4 weeks before being randomized into treatment arms (18-26 mice/ arm) and initiated on treatment. Untreated mice were sacrificed (i) the day after each infection to quantify CFU implanted in the lungs, and (ii) at initiation of treatment to determine pretreatment CFU counts.

#### H37Rv strain

Frozen and titrated *M. tuberculosis* H37Rv stock was diluted at 1:30 by 7H9 media with Tween 80 and used for infection, with the goal of implanting approximately 2.5 log_10_ CFU. Treatment started 4 weeks after infection. Untreated mice were sacrificed same as in the HN878 study.

### Chemotherapy

Rifampin, isoniazid, pyrazinamide and ethambutol were obtained from Sigma and formulated in distilled water. Pretomanid, linezolid, bedaquiline were provided by the Global Alliance for TB Drug Development. Pretomanid was prepared in the CM-2 (cyclodextrin micelle) formulation, linezolid in 0.5% methylcellulose, and bedaquiline in an acidified 20% hydroxypropyl-β-cyclodextrin solution, as previously described (28, 43). Dosing formulations were prepared weekly and stored at 4°C. All drugs except linezolid were administered once daily 5 days per week (5/7). In Experiment 1, linezolid was administered 3 or 6 days per week (3/7 or 6/7). In Experiments 2 and 4, linezolid was administered 5 days per week (5/7). In Experiment 3, linezolid was administered 3 or 5 days per week (3/7 or 5/7). All drugs were given by gavage. Pretomanid and bedaquiline were given at least 4 hours before linezolid (44). RHZE was given once daily with rifampin administered 1 hour prior to HZE. Doses used were as follows: bedaquiline 25 mg/kg, pretomanid 50 or 100 mg/kg (as indicated), rifampin 10 mg/kg, isoniazid 10 mg/kg, pyrazinamide 150 mg/kg and ethambutol 100 mg/kg. Linezolid doses were 45 or 90 mg/kg, or, in Experiment 2, were 25, 50 or 100 mg/kg (all as indicated).

### Assessment of treatment efficacy

Treatment efficacy was assessed using lung CFU counts at 1, 2 and/or 3 months of treatment and, in Experiment 3, the proportion of mice with culture-positive relapse, defined as ≥ 1 CFU detected after plating the entire lung homogenate obtained 3 months after treatment completion. CFU counts were determined by plating neat samples and serial 10-fold dilutions of lung homogenates on OADC-enriched selective 7H11 agar (Difco) supplemented with 0.4% activated charcoal as previously described (45, 46) to reduce carryover effects. Plates were incubated for at least 6 weeks before final CFU counts were obtained. In Experiment 4, the selection of drug-resistant bacteria after 2 months of treatment was determined by plating aliquots of the lung homogenates in parallel on 7H11 agar with or without bedaquiline 0.06 μg/mL or pretomanid 2 μg/mL to determine the proportion of the total CFU recovered that was resistant to either drug.

### Assessment of treatment toxicity

Blood was collected by cardiac puncture from a separate cohort of uninfected mice treated for 2 months alongside infected mice in Experiment 4. Complete blood counts (CBC) were performed in the Johns Hopkins Phenotyping Core, as previously described (13).

### Statistical analysis

Mouse lung CFU counts were log transformed before analysis. In Experiment 1 median CFU counts for BPa and BPaL groups were compared using a nonparametric t-test with Mann-Whitney post-test. In Experiment 2 mean CFU counts from the different treatment arms were compared using nonparametric analysis with Kruskal-Wallis post-test. In Experiments 3 and 4 mean CFU counts from the different treatment arms were compared using 1-way ANOVA with Dunnett’s post-test. Group proportions of mice with drug-resistant CFU were compared using Fisher’s Exact test. Group means for CBC parameters were compared using the same test. Analyses were performed using GraphPad Prism, version 6 (GraphPad, San Diego, CA).

### *In vitro* checkerboard assay

*M. tuberculosis* HN878 and H37Rv were grown in complete 7H9 media with Tween 80 then diluted with 7H9 media without Tween 80 until a turbidity of OD_600_=1.0 was reached. 100 uL of bacterial suspension was added to each well of 96-well polystyrene plates. Serial 2-fold dilutions were made for both drugs, starting at 4 μg/mL for linezolid and 0.96 μg/mL for bedaquiline. 50 μL each drug concentration was added per well to reach a final volume of 200 μL. Three studies were performed, each of which was 14 days in duration. The first study readout was turbidity. The next two studies used the Alamar blue assay. After 13 days of incubation, 25 μL of 0.02% resazurin solution was added per well and the color was recorded 24 hours later. The MIC was defined as the lowest concentration at which a color change from blue to pink was not seen. Assessment of drug interactions was based on calculation of the FICI values using the following equation:

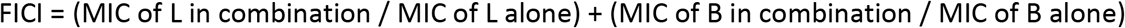

Results for each interaction were defined as synergistic if FICI values were ≤ 0.5, indifferent if FICI value was between 0.5 and 4, antagonistic if FICI value was > 4 (27).

## ACKNOWLEDGMENTS

This work was supported by the National Institutes of Health (R01-AI-111992, U19-AI-142735) and by the Global Alliance for TB Drug Development. KED is supported by K24AI50349. The authors would also like to thank Si-Yang Li for his help with the mouse model experiments.

